# Anterior Gradient-2 (AGR2) is overexpressed in colon cancer and is a potential biomarker of microsatellite instability (MSI) tumors

**DOI:** 10.1101/2021.09.07.459258

**Authors:** Delphine Fessart, Isabelle Mahouche, Veronique Brouste, Valerie Velasco, Isabelle Soubeyran, Pierre Soubeyran, Simon Pernot, Eric Chevet, Serge Evrard, Jacques Robert, Frederic Delom

## Abstract

**Background:** Colon cancer is one of the most common leading causes of death worldwide. Prognostic at an early stage is an efficient way to decrease mortality. The Endoplasmic Reticulum (ER)-resident protein anterior gradient-2 (AGR2), a Protein Disulfide Isomerase (PDI) is highly expressed in various tumours and is involved in tumour-associated processes. This study aims at examining the expression of AGR2 protein in colon cancer.

**Methods:** AGR2 protein expression was determined using immunohistochemistry on tissue samples issued from a cohort of 82 colorectal carcinomas.

**Results:** AGR2 protein expression was significantly higher in tumours than in adjacent nontumour controls. AGR2 expression subgroup analyses indicated that AGR2 low expression in colon cancer patients was significantly associated with worse overall survival. Mucinous colon cancers exhibited higher AGR2 expression levels than non-mucinous cancers. Additionally, tumours with microsatellite instability (MSI) were characterised by a strong upregulation of AGR2 mRNA and protein expression despite an absence of MLH1/MSH2 mutations.

**Conclusions:** Our findings indicate that high AGR2 protein expression is correlated with longer patient survival and that AGR2 overexpression is associated with MSI tumours and could represent an MSI biomarker. Overall, AGR2 might serve as a biomarker to stratify colon tumours and to contribute to the prognosis of colon cancer patients.

## 1 Bacground

Colon cancer, with 1.2 million new patients diagnosed worldwide each year, is the third most common malignancy [1]. Patient survival largely depends on the disease stage at diagnosis. Indeed, although this disease is potentially curable at early stages, the tumour frequently does not become symptomatic before advanced stages and is accompanied by malignant proliferation, extensive invasion, lymph node and distant metastasis [2]. Despite the advances that have been made by surgical techniques and chemotherapeutic options, 20% of patients die from recurrence and metastasis, accounting for poor prognostic outcomes [1]. Thus, there is a need for novel therapeutic targets and prognostic factors to further intensify and individualise patient care.

Most of tumour cells can control their microenvironment through the biological material they release. This control of tumour microenvironment occurs in part via proteins secreted by tumour cells, namely the tumour secretome. This implies that tumour cells, which are known to present alterations of protein homeostasis, can take advantage of this property to adjust the capacity of their secretory pathway to the protein folding and secretion demand. Since homeostasis of the secretory pathway in cancer is known to be mainly controlled by the first compartment of this machinery, the endoplasmic reticulum (ER) [3], we aimed at identifying ER molecular components that could affect the control of the tumour cell secretome. To this end, we undertook a proteomic analysis of the ER machinery [4], which led to the identification of a protein named Anterior Gradient 2 (AGR2 or PDIA17).

AGR2 is an ER-resident protein belonging to the PDI protein superfamily. In addition, we showed that AGR2 expression is controlled by the unfolded protein response (UPR) upon ER stress and most likely depends on both IRE1 and ATF6 signalling [4]. AGR2 expression is often upregulated in diverse epithelial cancers, allowing tumour cells to cope with abnormal protein production and contributing to the aggressiveness of cancer [5–7]. Its presence in biological fluids (blood, urine, serum) has been reported, showing that cancer cells secrete AGR2 into the extracellular space [5–7]. It has been demonstrated that other PDIs are also found in the extracellular milieu. For instance, we have shown that PDIA2 is secreted into the lumen of the thyroid follicles by thyrocytes to control extracellular thyroglobulin folding and multimerisation [8, 9]. Recently, we spotlighted extracellular AGR2 (eAGR2) as an inducer of Epithelial-to-Mesenchymal Transition (EMT) [7] and demonstrated that eAGR2 acquires extracellular gain-of-functions in the remodelling of epithelial tissue architecture, and exerts autocrine/paracrine signalling on stromal and cancer cells [5, 7], We also revealed that eAGR2 was able to selectively promote monocyte attraction, thereby linking eAGR2 to pro-inflammatory phenotypes [10]. At last, we recently showed that, in cancer cells, AGR2 was constitutively refluxed from the ER lumen to the cytosol (cytosolic AGR2 (cAGR2)) where it impedes the functions of tumour suppressors [11]. AGR2 has different roles in tumour-associated processes such as differentiation, proliferation, migration, invasion and metastasis [4–7, 12]. It has been shown that AGR2 protein overexpression may predict worse patient outcome in solid tumours [5–7, 13]. However, in a colon cohort published by Riener *et al*. [14], AGR2 protein expression appeared lost or decreased in the majority of colon tumours and the loss of AGR2 expression was significantly associated with reduced overall survival time.

To further clarify the relationships between AGR2 expression and colon cancer, we investigated its protein expression in tumour and adjacent non-tumour colon tissue samples, in a retrospective cohort of eighty-two primary colorectal carcinoma patients, as well as its mRNA expression from a public colon cancer database (The Cancer Genome Atlas (TCGA)), and correlated our findings to overall survival in univariate and multivariate analyses.

## 2 Materials and methods

### 2.1. Human tumour samples

The study was approved by the institutional review board of Institut Bergonié national ethics committee (NFS96900 Certification) and was performed in accordance with the Good Clinical Practice guidelines of the International Conference on Harmonization. Written informed consents were obtained from all patients. All medical records were de-identified. Samples were processed for histology studies.

### 2.2. Cohort

Eighty-two patients with colorectal carcinoma at different stages of evolution, operated and treated at Institut Bergonié, Bordeaux, France, between 1997 and 2005, were included in the study (Table 1).

**Table 1:** Patients Characteristics

| Variable | No. of patients (%) |  |  | P-Value |
| --- | --- | --- | --- | --- |
|  | Patients | AGR2 low (%) | AGR2 high (%) |  |
| **All cases** | 82 | 26 (31.7) | 56 (68.3) |  |
| **Patient age\*** |  |  |  |  |
| ≤65 years | 41 (50.0) | 17 (65.4) | 24 (42.9) | 0.06 |
| >65 years | 41 (50.0) | 9 (34.6) | 32 (57.1) |  |
| **pT-Status\*** |  |  |  |  |
| T1/T2 | 16 (19.5) | 3 (11.5) | 13 (23.2) | 0.03 |
| T3 | 46 (56.1) | 11 (42.3) | 35 (62.5) |  |
| T4 | 18 (22.0) | 10 (38.5) | 8 (14.3) |  |
| TX | 2 (2.4) | 2 (7.7) |  |  |
| **pN-Status\*** |  |  |  |  |
| N0 | 37 (45.1) | 3 (11.5) | 34 (60.7) | <0.001 |
| N+ | 43 (52.5) | 21 (80.8) | 22 (39.3) |  |
| NX | 2 (2.4) | 2 (7.7) |  |  |
| **Metastasis** |  |  |  |  |
| M0 | 54 (65.9) | 14 (53.9) | 40 (71.4) | 0.12 |
| M1 | 28 (34.1) | 12 (46.1) | 16 (28.6) |  |

\* Numbers may vary, since the information was not available for all patients.

### 2.3. Immunochemistry

Tissue fragments of the primary colon cancer and adjacent non-tumour were obtained after surgery and were fixed in Holland Bouin’s solution and paraffin-embedded for pathological evaluation. These samples were reviewed by an expert pathologist in the field (I. Soubeyran) and were used for the quantification of AGR2 protein expression by immunohistochemistry. Immunostaining was done on paraffin-embedded Tissue MicroArray sections (4 μm). In all available cases staining was performed on the Benchmark ultra-automated Stainer (Ventana) using diamino-benzidine as chromogen (DAB detection kit; Ventana, Roche), and anti-AGR2 monoclonal antibody (1:100; Abnova). A positive control (lung adenocarcinoma cell line A549 from ATCC) was tested on each immuno-histochemical run. Tissue sections were counterstained with hematoxylin and eosin stain (H&E). Images of H&E and AGR2 were acquired using a Leica DMIL LED microscope equipped with a Plan Fluor 10× 0.3 NA objective, and images were acquired using LAS X imaging software (Leica). Assessment of AGR2 immuno-histochemical staining was performed by both visual counting and image analysis using ImmunoRatio^®^. The cell counting tool selected for AGR2 indexing was the automated cell counter for digital histology slides ImmunoRatio (a free web application) [15], as it is available as a free tool with both browser-based analysis and an installable plugin for the open-source ImageJ (NIH, Bethesda, Maryland) image analysis software and does not require a high-end computer configuration [15]. Quantification was done by analyzing 8 images per sample.

### 2.4. Data source

The PanCancer Atlas (colorectal adenocarcinomas) of the Cancer Genome Atlas (TCGA) was used as the source of information for RNA-seq transcriptome data and clinical information. They were downloaded via the cbioportal (https://www.cbioportal.org).

### 2.5. Statistical analysis

The significance of associations between AGR2 expression and tumour characteristics was evaluated by Chi-2 test. Differences in expression of AGR2 in non-tumour and tumour tissue were evaluated using the Wilcoxon test on paired data. Univariate analysis of patients’ overall survival was carried out with the Kaplan-Meier method; all causes of death were considered as events, with a delay calculated between diagnosis and death. Patients living or lost to follow-up were censored with a participation time ranging from diagnosis to the last consultation or latest news. The log-rank test was used for comparison of tumour characteristics, variables significant at the 0.05 level in the univariate analysis were introduced in the multivariate model, which used a Cox stepwise descending maximum likelihood method.

## 3 Results

### 3.1. AGR2 immunostaining in colon cancer

Immunohistochemical (IHC) staining with anti-AGR2 antibodies was performed on 82 tissue samples from the cohort (Table 1) at Bergonie Institute (Bordeaux, France). The population of 82 patients, recruited over 1997–2005, consists of 40 women (48.8%) and 42 men (51.2%) (Table 1). The median age was 65 years (Table 1). Fifty-four patients (65.9%) were not metastatic and 28 patients (34.1%) were metastatic at diagnosis. We observed that AGR2 expression was significantly higher in Tumour tissues (T) compared to Adjacent Non-Tumour tissues (ANT) (Figure 1A) (p⍰0.001). The patient cohort was stratified in three groups, as having weak (⍰20%), moderate (20-49.9%) and strong (≥ 50%) AGR2 protein level expression (T tissues compared ANT tissues) (Table 2). None of the tumours was devoid of AGR2 expression, 26 patients (31.7%) exhibited AGR2 weak expression, 32 patients (39%) exhibited AGR2 moderate expression and 24 patients (29.3%) exhibited AGR2 strong expression. Representative images of AGR2 IHC staining of these tumour groups are shown in Figure 1B, C and D. These results show a clear association between high AGR2 expression and tumour tissue.

**Figure 1.**
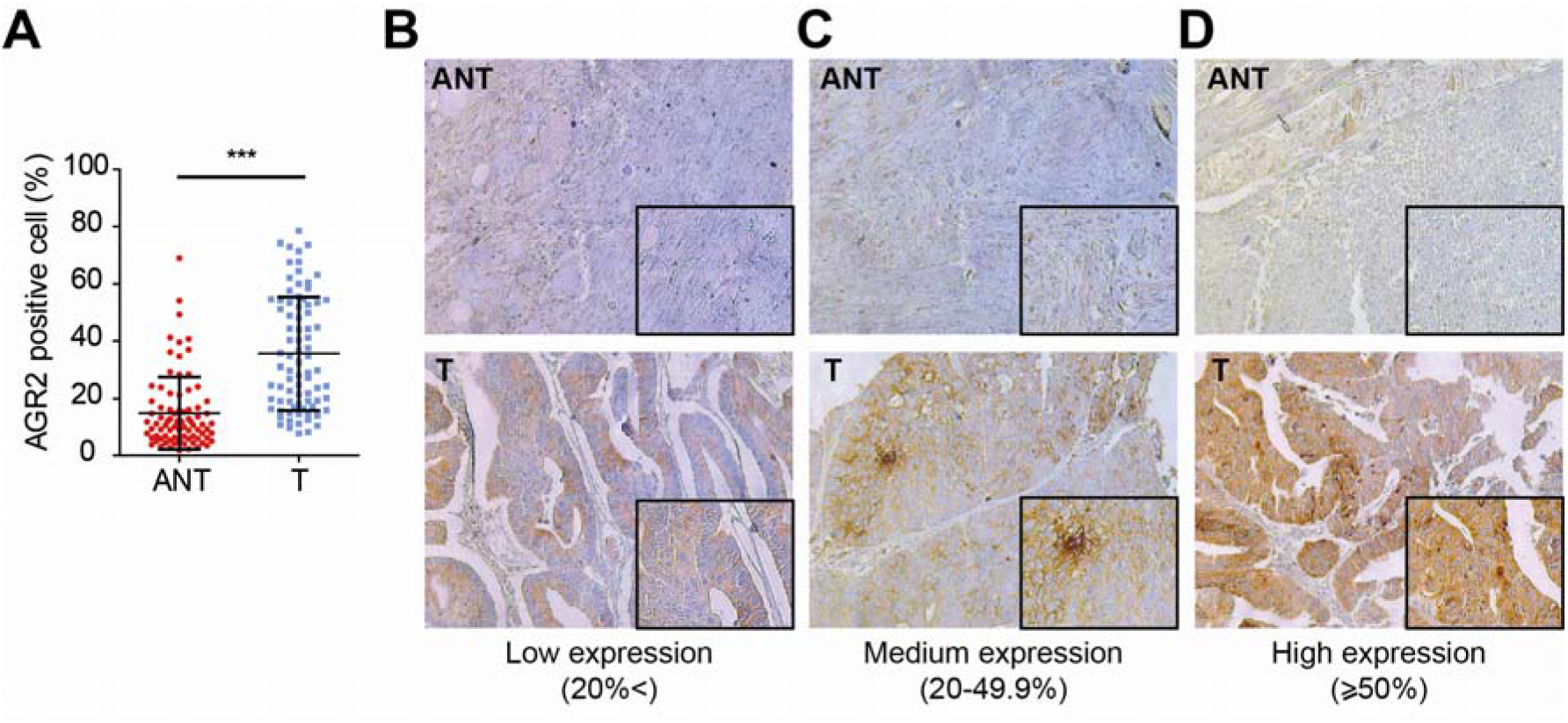
AGR2 immunohistochemistry expression in colon cancer tissue. **(A)** Box plot of the statistical analysis of the percentage of AGR2 positive cells between tumour (T) and corresponding Adjacent Non-Tumour (ANT) tissues. ***p<0.001 (Wilcoxon test). (B-D) AGR2 expression determined by immunohistochemistry in sections of fixed-paraffin-embedded colon tumour tissues (T) from patients at different stages as compared to Adjacent Non-Tumourtissues (ANT) (x100).

**Table 2:**
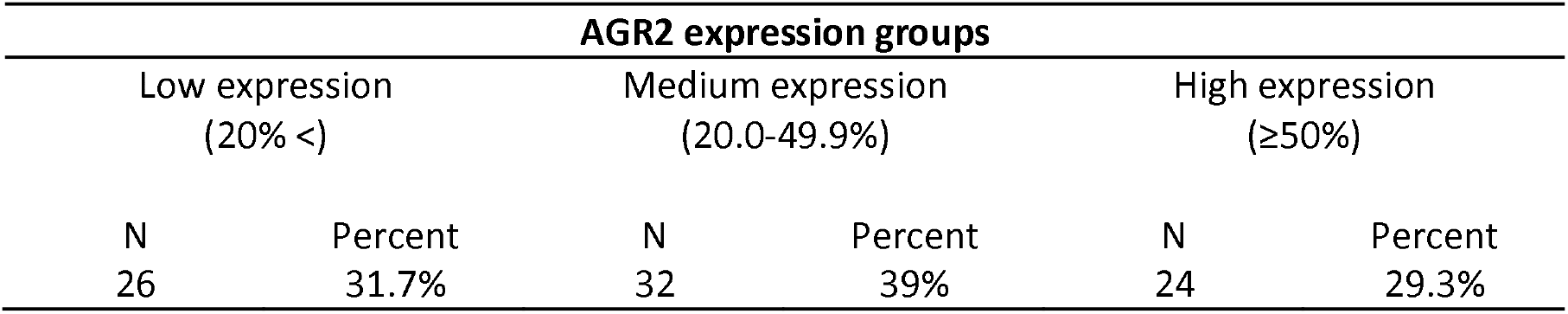
Stratification in three groups of AGR2 expression levels

| AGR2 expression groups |  |  |  |  |  |
| --- | --- | --- | --- | --- | --- |
| Low expression<br>(20% <) |  | Medium expression<br>(20.0-49.9%) |  | High expression<br>(≥50%) |  |
| N | Percent | N | Percent | N | Percent |
| 26 | 31.7% | 32 | 39% | 24 | 29.3% |

### 3.2. Association of AGR2 expression with clinical-pathological parameters

For statistical reasons, tumours were grouped according to AGR2 expression level either as low (31.7%, negative or weak) or high (68.3%, moderate or strong) (Table 1). AGR2 expression showed a significant association with Nodal status (pN-status (p⍰0.001, Table 1)) and Tumour status (pT status (p=0.03, Table 1)), high AGR2 expression being associated with both lower pT-status and lower pN status. No significant associations of AGR2 expression with patient age and metastatic status were found (Table 1).

### 3.3. Univariate and multivariate survival analyses

To test whether the changes in AGR2 expression were associated with colon cancer patient outcome, we performed univariate and multivariate survival analyses. As expected, pT-stage (Figure 2A), nodal status (Figure 2B) and metastasis status (Figure 2C) were significantly associated with reduced patient overall survival in univariate analyses (Table 3). It is remarkable that higher AGR2 expression was also significantly associated with longer patient survival (Table 3, Figure 2D). In multivariate overall survival analysis (Table 3), as expected, metastasis was an independent prognostic factor for overall survival (Table 3) and the combined HR for low *vs*. high AGR2 expression for overall survival was 0.35 (95% CI, 0.19 to 0.64; p=0.0006; Table 3). Thus, AGR2 expression was also an independent prognostic factor for overall survival in this multivariate analysis (Table 3).

**Figure 2.**
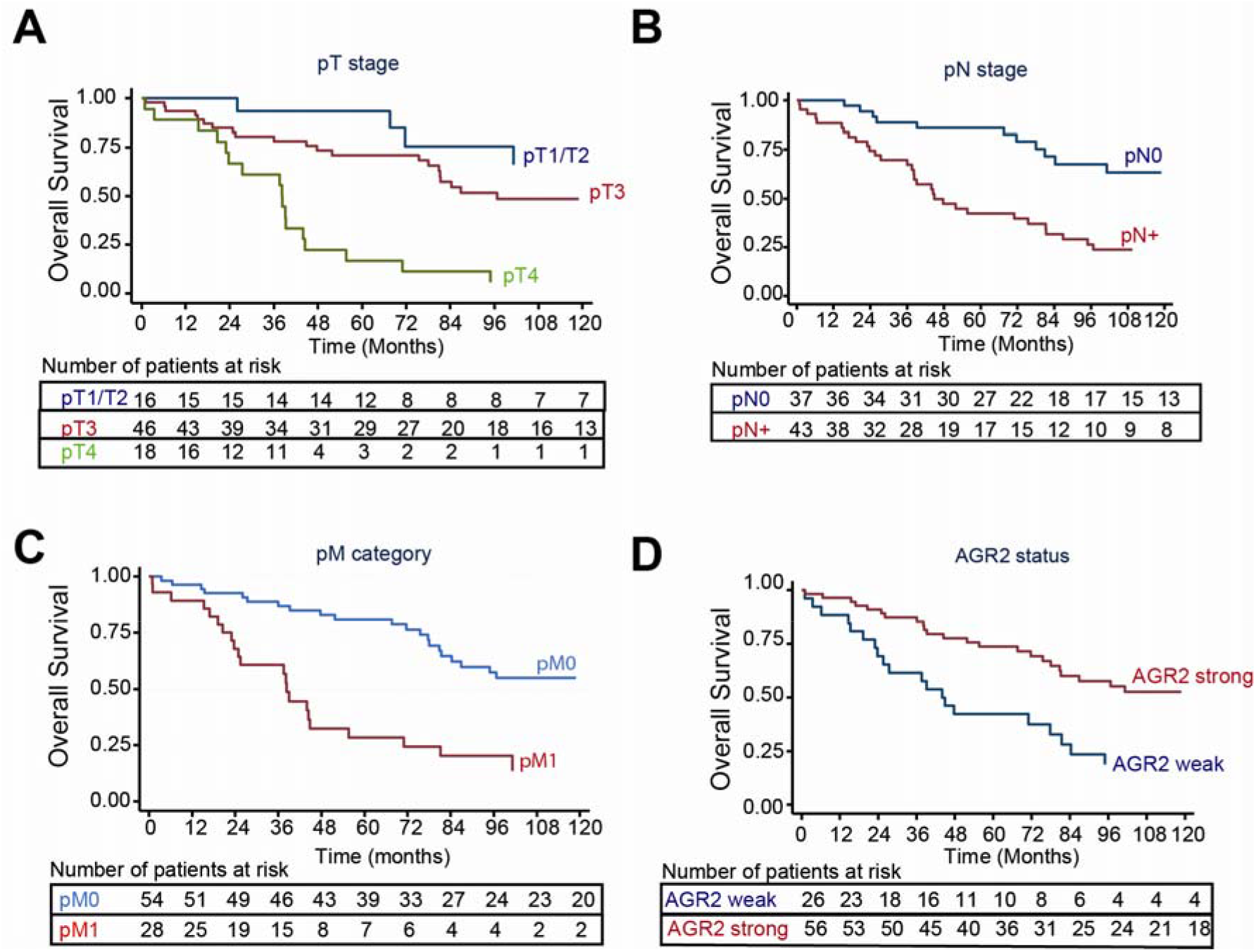
Kaplan–Meier curve for overall survival in colon cancer. **(A)** Tumour status (pT stage). **(B)** Nodal status (pN stage). **(C)** Metastasis status (pM category). **(D)** AGR2 protein expression status (AGR2 status).

**Table 3:**
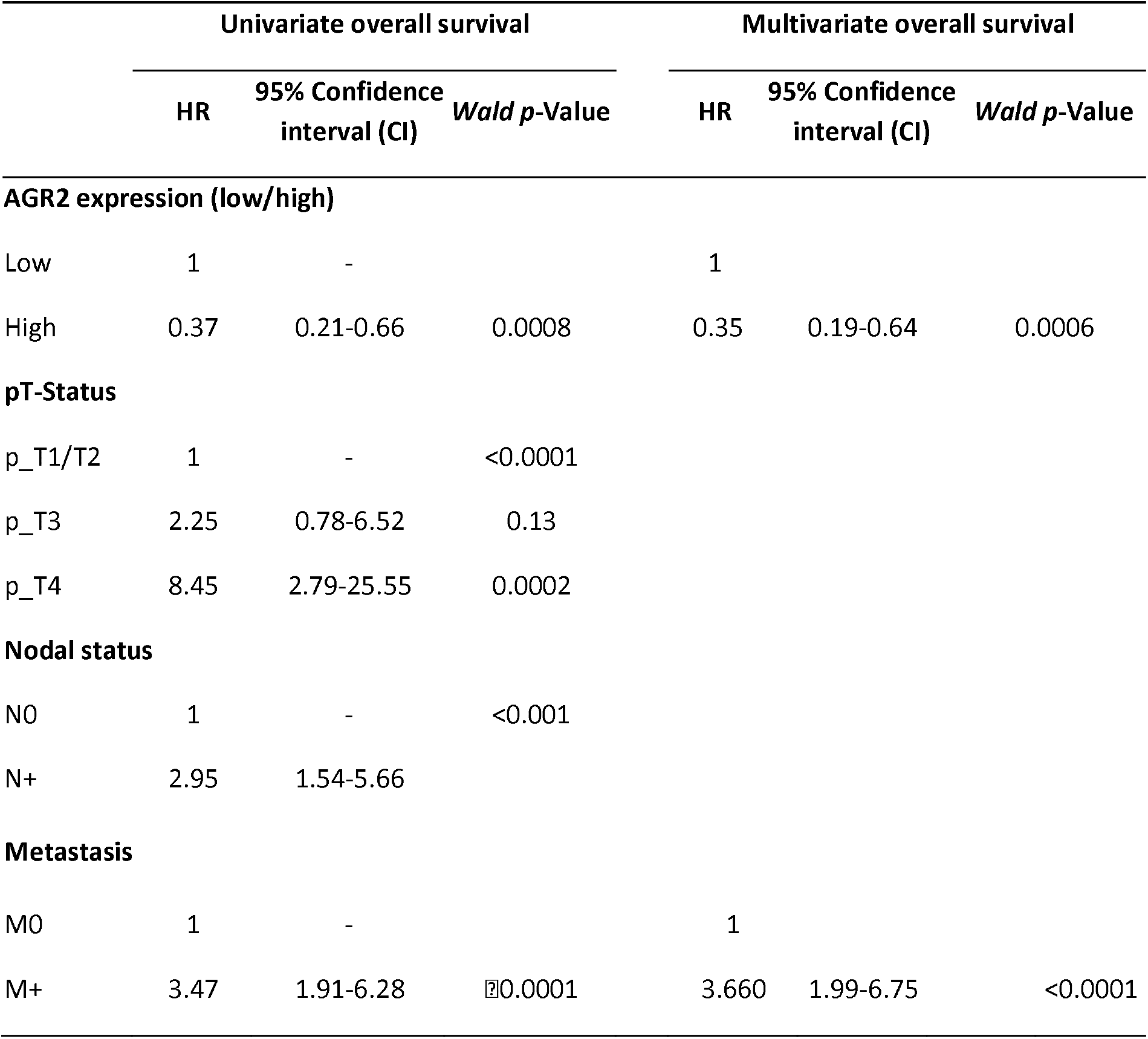
Influence of clinical-pathological parameters and AGR2 expression on patient survival

### 3.3. TP53 mutation and AGR2 expression

*TP53* mutation occurs in approximately 40%-50% of sporadic colon cancer [16] and is closely related to the progression and outcome of sporadic CRC. It has been reported that AGR2 expression could be linked to the *TP53* status [17]. In our cohort, we observed a decreased of AGR2 expression in TP53-mutated sample (Figure 3A). This was further confirmed in the analysis of AGR2 expression in colon cancer samples from TCGA datasets that AGR2 displayed significantly lower expression in TP53-mutated as compared to *TP53*-wild type colon cancer patients (Figure 3B). Taken together, these results show a strong association between AGR2 expression levels and *TP53* status.

**Figure 3.**
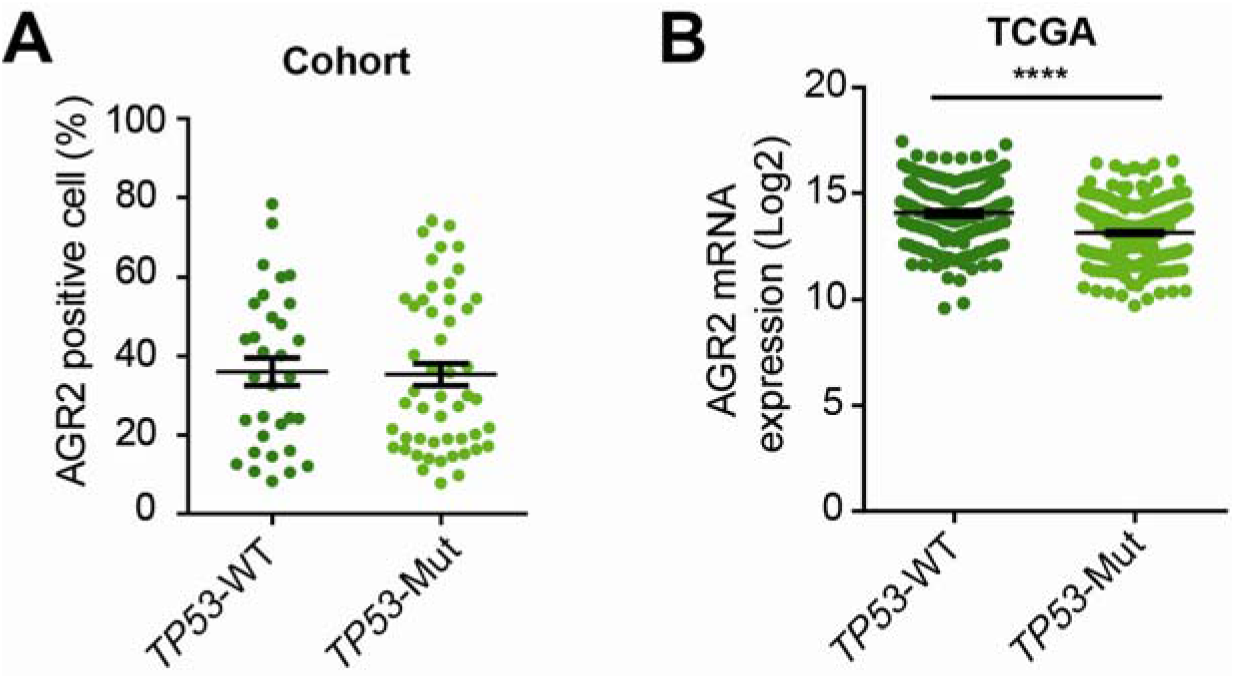
AGR2 expression is associated with TP53 status. **(A)** Percentage of AGR2 positive cells in TP53-wild type (*TP53*-WT) and TP53-mutated (TP53-Mut) tumours in our cohort study. **(B)** AGR2 expression in *TP53*-WT and TP53-Mut colon cancer tumour samples in TCGA database (p <0.0001).

### 3.5. AGR2 is upregulated in mucinous colon cancer

AGR2 expression has often been associated with mucinous tumours [18]. Indeed, a major characteristic of AGR2 expression is to be strongly overexpressed in mucin-secreting cells [19] and in mucinous adenocarcinomas subtypes [13]. In our cohort, the trend was an increase of AGR2 expression in mucinous tumours (n=4) as compared to non-mucinous tumours (n=78) (Figure 4A). The analysis of AGR2 expression in colon cancer samples from TCGA datasets showed that AGR2 displayed significantly higher expression in mucinous compared to non-mucinous tumours (Figure 4B). Taken together, these results show a correlation between high AGR2 expression levels and the mucinous colon cancer subtype.

**Figure 4.**
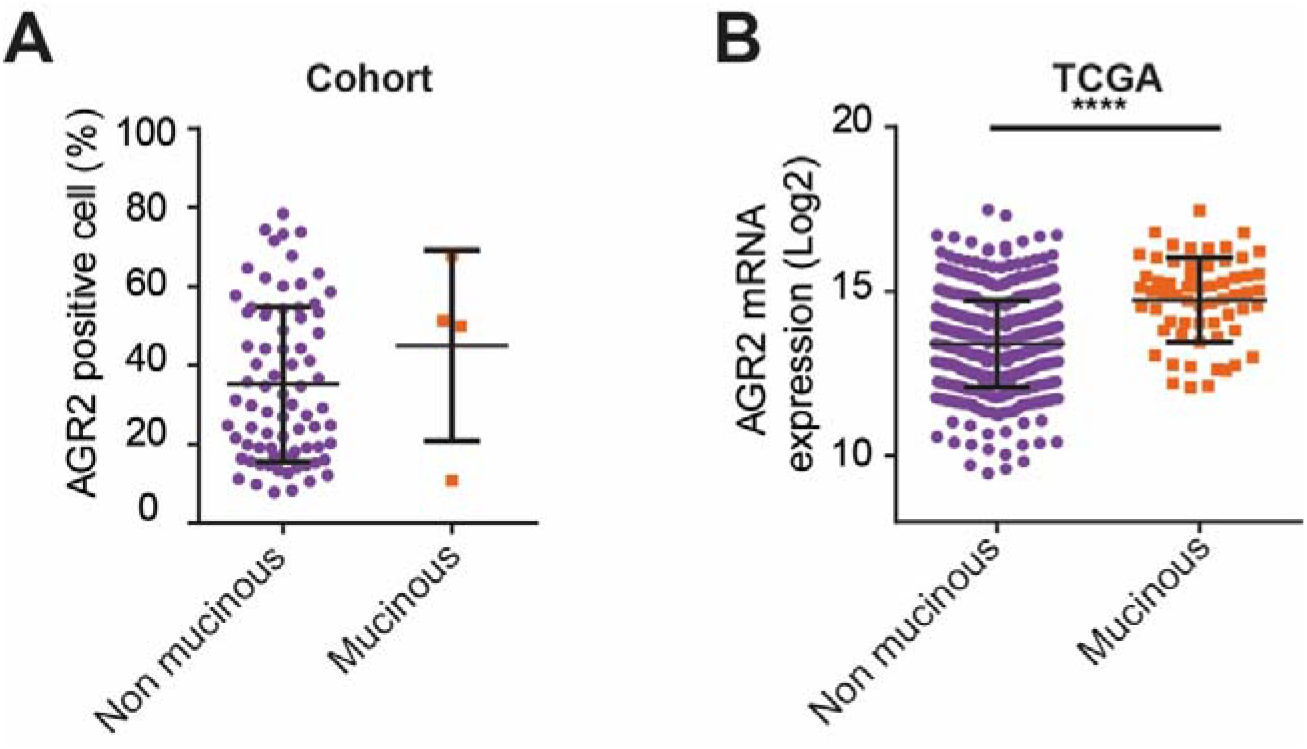
AGR2 expression is associated with mucinous colon cancer type. **(A)** Percentage of AGR2 positive cells in non-mucinous and mucinous tumours in our cohort study. (B) AGR2 expression in 531 non-mucinous and 61 mucinous colon cancer tumour samples in TCGA database (p = 2.5 × 10^-13^).

### 3.6. Microsatellite instability (MSI) tumours exhibit higher AGR2 expression

The MSI phenotype corresponds to a deficiency in mismatch repair (MMR) occurring in about 15% of colorectal tumours, while most tumours display proficient MMR status [20–22], In our patient cohort, *AGR2* expression was significantly higher in MSI (n=9) than in non-MSI tumours (n=73) (Figure 5A). Colorectal tumours from the TCGA database (459 samples) are divided in four subtypes according to classical criteria [22, 23]: 1) the microsatellite instability (MSI) group accounts for 13.7% of tumours; 2) the chromosome instable (CIN) group, which presents major chromosomal instability and accounts for 71.5% of tumours; **3)** the genome stable (GS) group (12.6%) and 4) the polymerase ε mutated (*POLE*) group (2.2%).

**Figure 5.**
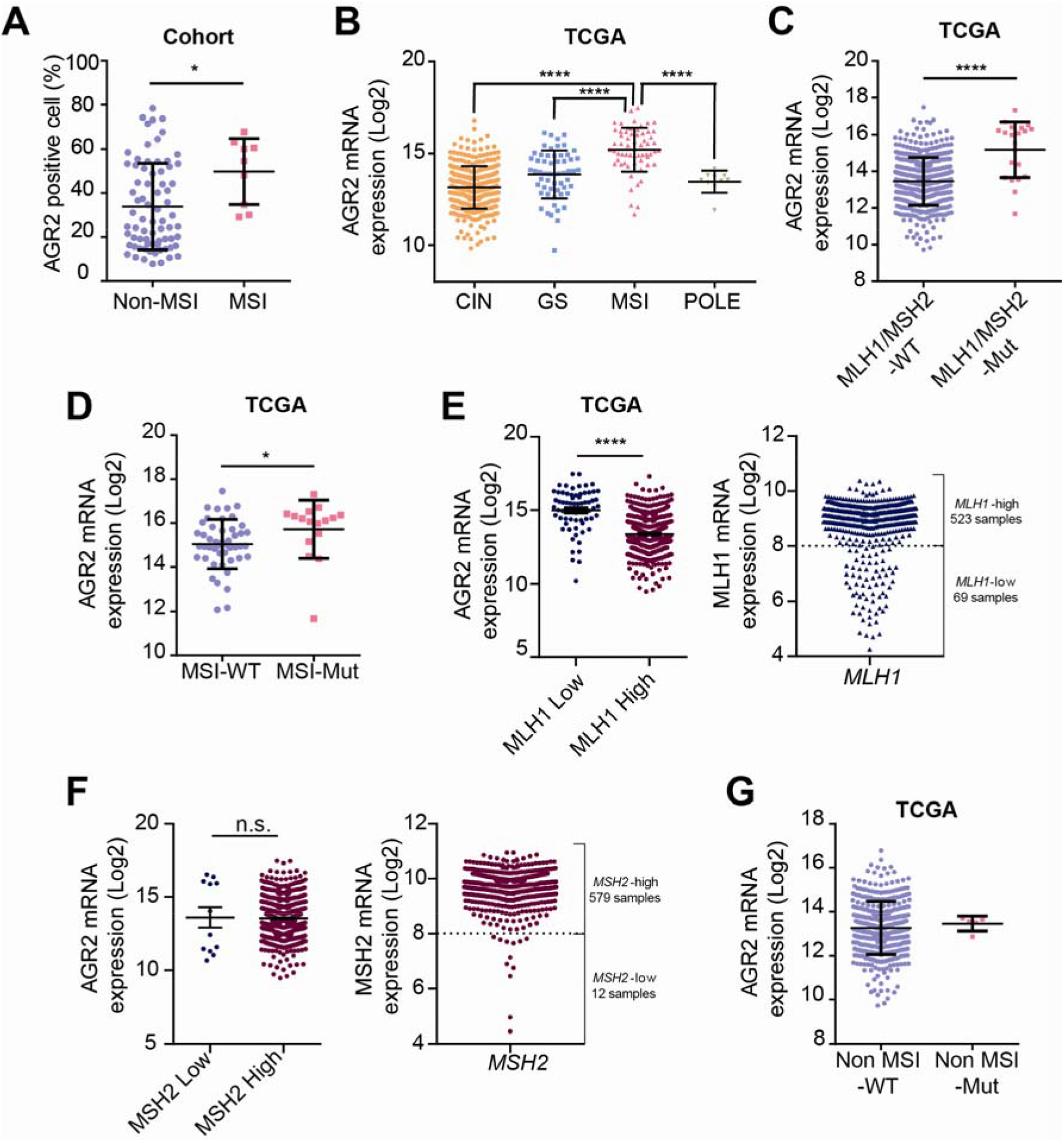
AGR2 expression in colon cancer based on the MSI status. **(A)** Percentage of AGR2 positive cells in non-MSI and MSI tumours in our cohort (p⍰=⍰0.0462). **(B)** *AGR2* expression in 459 colon tumour samples from TCGA database distributed according to tumour subtype (CIN: chromosome instable, 328 samples; GS: genome stable, 58 samples; MSI: microsatellite instable, 63 samples; POLE: polymerase ε mutated, 10 samples). AGR2 expression in MSI samples was different from its expression in CIN (p = 1.1 × 10^-31^), GS (p = 3.0 × 10^-8^), and POLE (p = 2.7 × 10^-5^), but was not significantly different in CIN, GS and POLE samples. **(C)** AGR2 expression in 466 tumour samples from TCGA database: comparison between MLH1/MSH2 mutated samples to MLH1/MSH2 wild-type samples (p = 9.24 × 10^-9^). (D) AGR2 expression in 63 MSI tumour samples from TCGA database: comparison according to the MLH1/MSH2 mutation status (p = 0.052). **(E) Left panel,** Differential *AGR2* expression in *MLH1*-low and *MLH1*-high colorectal cancer samples from TCGA. **Right panel.** The threshold between MLHl-low and MLHl-high samples was determined graphically from data plotting. **(F) Left panel,** Differential *AGR2* expression in *MSH2*-low and *MSH2*-high colorectal cancer samples from TCGA. **Right panel.** The threshold between MSH2-low and MSH2-high samples was determined graphically from data plotting. **(G)** AGR2 expression in 396 non-MSI tumour samples from TCGA database: comparison according to the MLH1/MSH2 mutation status (the difference is not significant).

The analysis of *AGR2* gene expression in colon cancer samples from the TCGA datasets supports our experimental results: *AGR2* displays a significantly higher expression in MSI tumours compared to all other tumour subtypes (Figure 5B). To understand the mechanism linking *AGR2* expression and MSI, we tried to associate AGR2 expression with the presence of mutations or reduced expression of *MLH1* and/or *MSH2*, the two main genes responsible for the MSI phenotype (generally by loss of expression in somatic MSI, and by deleterious mutations in MSI in hereditary predisposition). There was a significant difference in *AGR2* gene expression when comparing *MLH1/MSH2* mutated and WT samples in the whole collection of tumours (p = 2.27×10^-8^) (Figure 5C). However, the significance was reduced when considering only the group of MSI tumours (p = 0.052)(Figure 5D), probably because of the other cause of MSI (loss of MLH1/MSH2 expression) in the MSI group. Indeed, there was a highly significant inverse correlation between the expression of *AGR2* and that of *MLH1* (r = −0.298, p < 10^-7^) which can be shown by comparing *AGR2* expression in *MLH1*-high and *MLH1*-low samples (Figure 5E), but not with that of *MSH2* (Figure 5F). In contrast, there was no significant difference in *AGR2* gene expression between *MLH1/MSH2* mutated and WT samples in the cohort of non-MSI tumours (Figure 5G).

Taken together, our results demonstrate that 1) AGR2 is up-regulated in MSI tumours vs. all other colorectal tumour subtypes, and 2) this overexpression is due to contributions of both *MLH1/MSH2* mutations and loss of *MLH1* expression.

## 4. Discussion

AGR2 is a member of the protein disulphide isomerase family residing in the ER of epithelial cells [4–7]. This protein plays an important role in maintaining intestinal homoeostasis [24]. AGR2 is generally present in mucus-secreting epithelial cells and is highly expressed in Paneth and goblet cells of the intestine, with the highest levels in the ileum and colon [25]. Herein, we observed that AGR2 was expressed at higher levels in mucinous tumours than in non-mucinous tumours, which agrees with previous studies in mucinous cancers [13]. In goblet cells, it was suggested that AGR2 forms mixed disulphide bonds with Mucin 2 (MUC2), allowing for its correct folding and secretion [24, 26]. We showed previously that the release of extracellular AGR2 (eAGR2) in the intestinal microenvironment yields pro-inflammatory phenotypes. Hence, these AGR2 data in IBD might apply to cancer biology, since proteostasis imbalance emerged as a major cancer hallmark, capable of driving tumour aggressiveness [27]. Indeed, control of AGR2 may be a factor in tumourigenesis. High AGR2 expression, as well as its secretion (eAGR2) into the tumour microenvironment, has been reported, by us and others, in many cancer types, and has been associated with pro-tumourigenic phenotypes and poor patient outcomes [5-7, 13]. Although Anterior Gradient proteins, including AGR2, were identified more than 15 years ago, their precise biological functions remain unclear [5, 6, 19, 28].

In our study, we found 1) that AGR2 was highly expressed in colon tumour tissue as compared to non-tumour tissue, and **2)** that high expression of AGR2 is associated with longer patient survival time, even in a multivariate Cox analysis. Similarly, Riener et al. have shown [14] that the loss of AGR2 expression was significantly associated with reduced overall survival time. In agreement to those clinical observations, Gupta *et al*. [29] have shown that, in a constitutive *Agr2* KO mouse model, the absence of AGR2 expression resulted in a significant increase in proliferation of gastric epithelial cells.

In different human adenocarcinoma cohorts, it has been shown that AGR2 was associated with the Epithelial-Mesenchymal Transition (EMT) phenotype [5, 6] and we have already shown [7] that eAGR2 is sufficient by itself i) to induce EMT and ii) to generate invasive structures in non-tumour epithelial organoids, thus indicating that eAGR2 is an important determinant of epithelial morphogenesis disruption, invasion and metastatic dissemination. Here, our results show a correlation between AGR2 expression levels and *TP53* status. Indeed, *AGR2* displayed significantly higher expression in TP53-wild type as compared to TP53-mutated colon cancers. Recently, we demonstrated that AGR2 is refluxed from the ER to the cytosol (cytosolic AGR2 (cAGR2)) in cancer cells and, in the latter compartment, acts, through a gain-of-cytosolic function, as an inhibitor of p53 tumour suppressor activity [11]. Thus, our results suggest that in the TP53-wild type colon cancer sub-population, AGR2 could be increased to be refluxed in the cytosol (cAGR2) to inhibit the p53 tumour suppressor activity and thus to favour tumour development. As a conclusion, both abnormal AGR2 localisations provide selective advantages to tumour through gain-of-extracellular or -cytosolic functions, as a pro-oncogene.

Recently, a systematic review and meta-analysis on the prognostic value of AGR2 expression in adenocarcinomas have shown a significant correlation between AGR2 expression and poor overall survival in almost all adenocarcinoma patients [13]. Thus, although we and others had demonstrated significant survival benefits of patients with AGR2 low-level tumours, the present results on colon cancer patients, in agreement to those presented by Riener *et al*. [14], are opposite to the general trend observed in adenocarcinomas. Hence, we may speculate that the function and the effect of AGR2 might have tumour suppressor properties in certain malignancies, and more particularly in colon cancer, which necessitates further functional studies. Thus, we hypothesise that the different impact of AGR2 expression in adenocarcinomas might be due to a distinct regulation in various cancer types. Future research should clarify whether AGR2 overexpression is involved in tumourigenesis and metastatic dissemination in colon cancer.

It may be important to emphasise the fact that a high expression of AGR2 is especially observed in MSI tumours, both in our cohort and in the TCGA database. It appears that this is not only dependent on the *MLH1/MSH2* mutational status, but also to the loss of *MLH1* expression. As a consequence, the overexpression of AGR2 in MSI tumours appears to be related to the MSI phenotype itself, whatever the mechanism of its acquisition (genomic or epigenomic). The good prognosis of AGR2-overexpressing colorectal tumours might be related to the overall better survival of MSI tumours-bearing patients, as long as they are not metastatic. The association between MSI tumour phenotype, *AGR2* overexpression and good prognosis has to be more closely studied in prospective cohorts of colorectal patients, and the mechanism(s) underlying this association has to be deciphered.

In summary, we found that AGR2 is increased in colon cancer cells as compared with adjacent tissue, and that the overexpression of AGR2 is an independent factor of poor prognosis for survival in this cancer type. In addition, AGR2 is upregulated in mucinous colon cancer and high overexpressed in MSI tumours. In conclusion, AGR2 expression could be useful as a biomarker to stratify colon cancer phenotypes and to contribute to the prognosis of colon cancer patients.

## Abbreviations

ANT: Adjacent Non-Tumour tissues
AGR2: Anterior Gradient-2
CIN: Chromosome Instable
cAGR2: cytosolic AGR2
ER: Endoplasmic Reticulum
EMT: Epithelial-to-Mesenchymal Transition
eAGR2: extracellular AGR2
GS: Genome Stable
IHC: Immunohistochemical
MSI: Microsatellite Instability
MMR: Mismatch Repair
MUC2: Mucin 2
pN-status: Nodal status
POLE: Polymerase e mutated
PDI: Protein Disulfide Isomerase
TCGA: The Cancer Genome Atlas
pT-status: Tumour status
T: Tumour tissues
UPR: Unfolded Protein Response

## Declarations

### Ethics approval and consent to participate

The study was approved by the institutional review board of Institut Bergonié national ethics committee (NFS96900 Certification) and was performed in accordance with the Good Clinical Practice guidelines of the International Conference on Harmonization. Written informed consents were obtained from all patients. All medical records were de-identified.

### Availability of data and materials

Not applicable

### Consent for publication

The authors declare that their participation in writing this review as well as its publication is voluntary.

### Authors’ contributions

JR and EC provided guidance throughout the preparation of the manuscript. DF, FD and JR performed literature search, the figures and wrote the manuscript. IM, VB and VV contributed to the acquisition of data and the analysis and interpretation of data. IS, PS, SP, JR and SE contributed to the clinical expert point of view and critically revised the manuscript. EC contributed to the scientific expert point of view and critically revised the manuscript. All authors read and approved the final manuscript.

## Acknowledgements

We gratefully acknowledge the members from ARTiSt group for their critical remarks.

